# Functional single-cell profiling identifies that exosomes are associated with increased immune cell infiltration in non-metastatic breast cancer

**DOI:** 10.1101/2020.12.14.422613

**Authors:** Mohsen Fathi, Robiya Joseph, Jay R T. Adolacion, Melisa Martinez-Paniagua, Xingyue An, Konrad Gabrusiewicz, Sendurai A. Mani, Navin Varadarajan

**Author notes:** To whom correspondence should be addressed: Navin Varadarajan, Department of Chemical and Biomolecular Engineering, University of Houston, Houston, Texas TX 77204, USA.

## Abstract

Exosomes mediate intercellular communication in health and disease. Conventional assays are limited in profiling exosomes secreted from large populations of cells and are unsuitable for studying the functional consequences of individual cells exhibiting varying propensity for exosome secretion. In cancer, since exosomes can support the development of the pre-metastatic niche, cells with varying abilities to secrete exosomes can directly impact tumorigenesis. Here, we developed a high throughput single-cell technique that enabled the mapping of exosome secretion dynamics. By utilizing clinically relevant models of breast cancer, we established that non-metastatic cancer cells secrete more exosomes than metastatic cancer cells. Single-cell RNA-sequencing confirmed that pathways related to exosome secretion were enriched in the non-metastatic cells compared to the metastatic cells. We established isogenic clonal cell lines from non-metastatic cells with differing propensities for exosome secretion and showed that exosome secretion is an inheritable property preserved during cell division. Combined in vitro and in vivo studies with these cell lines suggested that exosome secretion can impede tumor formation. In human non-metastatic breast tumors, tumors with higher secretion of exosomes have a better prognosis, higher immune cytolytic activity, and enrichment of pro-inflammatory macrophages compared to tumors with lower secretion of exosomes. Our single-cell methodology can become an essential tool that enables the direct integration of exosome secretion with multiple cellular functions.

## Introduction

Exosomes, a subset of extracellular vesicles (EV), comprise a fundamental mechanism of intercellular communication across distant cells and serve to transport biological molecules such as lipid, nucleic acids, and proteins. Encapsulation of molecules into exosomes fundamentally alters their stability, transport, and trafficking, and characterizing the secretion of exosomal cargo from cells is of great interest in fundamental cell biology and for targeted drug delivery.^1-6^

In the context of cancer, exosomes are known to affect a variety of biological events that promote tumor progression such as angiogenesis,^7, 8^ invasion,^9^ evasion of immune surveillance,^10, 11^ and drug resistance.^12^ Exosomes from highly metastatic melanoma tumors promoted vascular permeability and contributed to the formation of the pre-metastatic niche.^13^ Also, exosomes can transfer antigens and enhance the immune response by activating T cells and NK cells.^14, 15^ Due to their stability, they have great potential in cancer diagnosis^16, 17^ and treatment.^18^ Mapping the dynamic secretion of exosomes at the cellular level can significantly advance our understanding of the role of exosomes in cancer.

From an analytical standpoint, the size of exosomes (40-150 nm) is in between the size of proteins and cells. A number of analytical methods including nanoparticle tracking analysis (NTA),^19^ electron microscopy,^20^ flow cytometry,^20, 21^ microfluidic devices,^22^ and western blotting^23^ have been widely used for characterization of exosomes, some even with the sensitivity of detecting individual exosomes. Unfortunately, however, these exosomes are derived from culturing billions of cancer cells and thus represent an averaging of exosomes secreted by all cells. From the standpoint of disease biology, this is suboptimal since these might reflect supra-physiological concentrations of these exosomes. Second, since tumors are heterogeneous populations, these approaches mask the inherent differences in exosome secretion between individual cells, and mapping the direct relationship between exosome secretion and tumorigenic potential is not feasible. Not surprisingly, recent advances in microfabrication have revealed that the rate of exosome secretion from single cells can be very different.^24-26^ Despite this progress, however, technological hurdles have prevented us from answering a number questions at the single-cell level including (1) heterogeneity of the short-term dynamics of secretion of exosomes, (2) whether exosome secretion is an inheritable property preserved upon cell division, and (3) whether there is a difference in tumorigenic potential between isogenic tumor cells with differences in the rate of exosome secretion.

Here we report a high-throughput single-cell technique for the dynamic quantification of exosome secretion from single cells. We utilized the 4T1 and 67NR syngeneic mouse mammary tumor models since these are well-validated, clinically relevant models with vastly different potential for metastasis. 4T1 spontaneously metastasizes to multiple sites, whereas 67NR is incapable of metastases and is restricted to the formation of the primary tumor.^27^ By tracking the dynamics of exosome secretion, we demonstrate that in both cell lines, the dominant secretor cells are capable of continuous secretion over short time intervals (6-24 hours). Surprisingly, the non-metastatic 67NR cells secreted more exosomes per cell than 4T1 cells, and this result was consistent with scRNA-seq of the same cells, showing an enrichment of the ALIX-Syndecan-Sytenin pathway. Although the secretion of exosomes from highly secreting 67NR clones caused an increase in proliferation and migration *in vitro*, the tumor growth was inhibited *in vivo*. Analysis of The Cancer Genome Atlas (TCGA) data illustrated that the secretion of exosomes is associated with better overall survival of non-metastatic patients, which was induced by higher secretion of IFN-γ, higher infiltration of Th1 cells, the polarization of M1 macrophages, and suppression of the IL6ST/STAT3 pathway. More broadly, the exosome secretion signatures are associated with better prognosis in non-metastatic melanoma but worse prognosis in non-metastatic lung cancers.

## Material and Methods

### Cell culture

4T1 and 67NR cells were purchased from ATCC. We cultured cells in RPMI 1640 supplemented with 10% FBS, 1% L-glutamine, HEPES, and penicillin-streptomycin. We cultured GSC20 cells in 50/50 mixture of Dulbecco’s Modified Eagle’s Medium (DMEM) and Ham’s F-12 medium supplemented with 1% penicillin-streptomycin, B-27 supplement, and epidermal growth factor. We tested all cells for mycoplasma contamination using real-time PCR.

### Exosome isolation and measurement

We used ultracentrifugation to isolate exosomes from GSC20 stem cells. Starting with 250 ml of culture media, we centrifuged the conditioned media at 300 × g for 4 minutes, filtered with 0.22 µm filters, and centrifuged at 10,000 × g for 30 minutes followed by ultracentrifugation at 100,000 × g for 70 minutes to pellet the exosomes. We washed the exosome pellet with PBS twice and centrifuged for another 100,000 × g for 70 minutes to purify the exosomes. We resuspended the exosomes in PBS and measured the exosome size distribution using the nanoparticle tracking analyzer (NTA). We stored the isolated exosomes at 4°C for one week or at -80°C for long term use.

### Bulk exosome detection assay

We performed immunoassays utilizing LumAvidin beads (Luminex, catalog number L100-L115-01) to capture exosomes with different protein markers. First, we centrifuged 10^5^ beads and resuspended in PBS with 1% BSA. We next incubated them with 3.5 µg/ml biotinylated CD81 or CD63 antibody (BioLegend, clone 5A6, and H5C6) for 30 minutes at room temperature, and washed them twice in PBS with 1% BSA. We added the exosomes at a 10^9^ particle/ml concentration and mixed on a rotator for 2 hours at room temperature, followed by washing in PBS with 1% BSA twice. We mixed the beads with 4 µg/ml PE anti-CD63 antibody (BioLegend, clone H5C6) and rotated for 45 minutes at room temperature. Finally, after two washes, we resuspended the pellet in PBS with 1% BSA and imaged using A1/TiE inverted confocal microscope (Nikon) equipped with 20x/0.75 NA objective. We measured the fluorescent intensity of CD63 on the beads using ImageJ.

### Functionalization of beads with anti-CD81 coating

We washed 10^5^ LumAvidin beads (Luminex, catalog number L100-L115-01) in PBS with 1% BSA and incubated the beads with 3.5 µg/ml biotinylated anti-CD81 antibody (BioLegend, clone Eat-2) at the room temperature for 40 minutes. Then, after washing beads thrice in PBS with 1% BSA, we resuspended them in 120 µl of PBS with 1% BSA.

### Exosome quantification using transwell assay

We utilized a Transwell insert with 3 µm pore membrane and loaded functionalized beads at the lower compartment, and cells on the upper compartment of the insert. For the GW4869 treatment assay, we used exosome-free complete media containing either 10 µM GW4869 or 10% DMSO. After 48 hours of incubation at 37°C, we collected the beads and labeled them with 4 µg/ml PE anti-CD63 antibody (BioLegend, clone NVG-2) for 45 minutes at 37°C. We subsequently washed the beads three times in PBS with 1% BSA and performed imaging using a Zeiss Axio Observer Z1 microscope equipped with 20x/0.8 NA objectives. Using ImageJ, we segmented and measured the fluorescent intensity of CD63 on the beads.

### Transmission Electron Microscopy (TEM)

Via exosome quantification using a transwell assay, and after 48 hours incubation at 37°C, we collected the beads and fixed with 2% glutaraldehyde (Ladd research, catalog number 20215). We placed the samples on 100-mesh carbon-coated, formvar-coated copper grids treated with poly-L-lysine for approximately 1 hour. We then negatively stained the samples with Millipore-filtered aqueous 1% uranyl acetate for 1 minute. The stain was blotted dry from the grids with filter paper, and samples were allowed to dry. We examined the samples in a JEM 1010 transmission electron microscope (JEOL, USA, Inc., Peabody, MA) at an accelerating voltage of 80 kV. We obtained the digital images were using the AMT Imaging System (Advanced Microscopy Techniques Corp., Danvers, MA).

### PDMS nanowell array fabrication and preparation

Applying standard soft lithography techniques, we fabricated the PDMS nanowell array as previously described.^28^ Before loading cells on the nanowell, we re-oxidized the array with air plasma and incubated with 1.5 ml PLL-g-PEG (SuSoS, Switzerland) solution dissolved in 10 mM HEPES buffer for 20 minutes at 37°C. After incubation, we rinsed the array with complete media before loading the cells.

### Single-cell exosome detection assay

To perform the single-cell assay for the detection of exosomes, we prepared the nanowell array and functionalized beads, as described above. We labeled 67NR or 4T1 cells with PKH67 dye (Sigma-Aldrich, catalog number PKH67GL-1KT) as directed by the manufacturer. We loaded labeled cells and functionalized beads, sequentially on the nanowell array. We covered the nanowell with exosome-free complete media and imaged at its initial time point, incubating at 37°C. Every two hours, we incubated the nanowell array with 4 µg/ml PE anti-CD63 antibody (BioLegend, clone NVG-2) for 45 minutes at 37°C. We subsequently washed the nanowell array three times in PBS with 1% BSA and performed imaging using microscopy. After each imaging, we returned the nanowell to the incubator at 37°C. We acquired all images by Zeiss Axio Observer Z1 microscope equipped with 20x/0.8 NA objectives and a Hamamatsu Orca Flash v2 camera.

### Secretion analysis of single-cell exosome detection assay

We analyzed the TIFF images from microscopy, as outlined in Figure S3. Briefly, for all images at each time point, we segmented the images into cells and beads and determined the ratio of the number of cells to the number of beads in each well. After identification of wells with a single bead and a single cell, we tracked the wells across time points. We background-corrected the CD63 pixel values and compared the pixel values between the bead and non-overlapped pixels on the cells using a two-tailed t-test. Based on the average intensity and the *p*-value calculated, we classified the single cells as either secretor (high secretor) or non-secretor (low secretor) cells.

### Kinetic analysis of single-cell exosome detection assay

Using the wells containing a single bead and a single cell identified in the secretion analysis of single-cell exosome detection assay, we selected the wells which were detected in all the time points for the kinetic analysis. To determine the behavior of the cells between time points, we performed a two-tailed t-test on the CD63 pixel values of the bead between two consecutive time points. We chose an increase in intensity with a *p*-value below 0.01 as the criterion for a significant change in the secretion behavior of the cell.

### Establishment of clonal cell lines

We retrieved the secretor and non-secretor single cells using a micromanipulator (ALS, CellCelector) equipped with 50 μm glass capillaries. We transferred single cells to a 96-well plate containing complete media. We monitored the single cells and cultured them in complete media until they proliferated to 24 population doublings.

### Wound healing assay

We cultured 67NR-S and 67NR-NS cells in a 12-well plate to 90% confluency with 10% FBS complete media. We subsequently replaced the media with 0.5% exosome-free FBS complete media for 12 hours. After starvation, we scratched the cells with 10 µl pipette tips and washed twice with PBS to remove the detached cells. We cultured the cells with 0.5% exosome-free FBS complete media during the assay to slow down cell proliferation. We obtained the images from six different areas per well with Zeiss Axio Observer Z1 microscope equipped with 20x/0.5 NA objectives at several time points. We analyzed the images with TScratch tool.^29^

### Soft agar colony formation assay

Performing an anchorage-independent growth assay using SeaPlaque agarose (Lonza, catalog number 50101), we assessed the transformation capacity of the 67NR-S and 67NR-NS cells *in vitro*. We used three different conditions in triplicates to determine the ability of these cells to form soft agar colonies: no treatment, 10% DMSO, and 10 µM GW4869 (Cayman Chemical, catalog number 13127). We suspended 2.5 × 10^3^ cells in 0.7% top agar in exosome-free complete media containing the appropriate treatment conditions and placed on top of solidified 0.8% bottom agar in 6-well plates (Fisher, catalog number 353046). Upon setting of the top agar with cells, we added 500 µl of fresh exosome-free complete media containing the appropriate treatment conditions to the wells and incubated the plates for 14 days at 37°C. We fed the cells with exosome-free complete media with the appropriate treatments, twice per week. We counted the colonies from ten different areas per well and acquired the representative 20x images microscopically using Zeiss Axio Observer A1 microscope.

### Mouse modeling assay

We injected 1 × 10^4^ 67NR-S and 67NR-NS cells subcutaneously into the fourth left mammary fat pad of five BALB/c mice (Jackson laboratory, strain 0000651 BALB/cJ) for each clone. We monitored the size of the tumor with caliper measurements weekly and calculated using formula (L × W^2^) × 0.5, where L and W are the length and the width of the tumor, respectively. We sacrificed the mice and harvested the tumors before the onset of necrosis.

### Single-cell RNA-sequencing

Following the Illumina Bio-Rad SureCell WTA 3’ library prep reference guide, we prepared the scRNA-seq library. Briefly, we mixed an equal number of the mouse cell lines in cold PBS with 0.1% BSA in a concentration of 2,500 cells/µl, then filtered to achieve single-cell suspension. Using a ddSEQ Single-Cell Isolator, we co-encapsulated with oil the single cells and barcodes into droplets. After reverse transcribing and breaking the emulsion, we purified the first-strand products using purification beads, followed by cDNA synthesis and tagmentation. We PCR-amplified the cDNA and cleaned it up to remove short library fragments. Later, we sequenced the cDNA library in a NextSeq 500 sequencing system. Using Illumina BaseSpace Sequence Hub, we analyzed the sequencing data and created a count matrix containing the number of transcriptomes for every single cell. We imported these matrices into R and combined them into a single matrix, which was then cleaned, normalized, and analyzed using the *Seurat* (v3.1.4) package.^30^ We ranked the differentially expressed genes of 4T1 and 67NR cell lines and transformed into human orthologous using the *BiomaRt* (v2.38.0) package,^31, 32^ and imported to GSEA software^33, 34^ provided by UC San Diego and Broad Institute for gene set enrichment analysis.

### Bulk RNA sequencing dataset analysis

We downloaded the raw counts of RNA-seq dataset published by Kim *et al*. from GEO (GSE104765).^35^ We filtered the table for three replicates of 4T1 and 67NR cells. To obtain the differentially expressed genes, we used the *DESeq2* (v1.22.2) package^36^ in R.

### Tumor Cancer Genome Atlas (TCGA) analysis

We downloaded all the TCGA data, including raw counts, RSEM gene normalized expression, and clinical data from the Broad Institute FireBrowse Data Portal (www.firebrowse.org). To collect the non-metastatic patients without lymph node metastasis, we used the TNM staging information and selected the patients with N0 and M0 for analysis. To perform hierarchal clustering on the breast cancer dataset, first, we filtered out genes with average RSEM expression < 5 to remove their effects in the data analysis. Next, we used *hclust* function in R to identify two clusters using *ward*.*D2* as the linkage method with *manhattan* as the distance measure. Using the *DESeq2* (v1.22.2) package, we identified CD63 and CD81 upregulated in cluster 1. Using a set of genes associated with exosome secretion (Table S1), we identified 13 genes with more than 1.2-fold change in cluster 1 as exosome signature genes for further analysis. For gene set enrichment analysis, we used the pre-ranked gene list of genes with a significant fold change of < 0.05 in GSEA software provided by UC San Diego and Broad Institute. For survival analysis, we used the Kaplan-Meier method to compare the overall survival of patients divided by the median expression of 13 exosome signature genes. We tested the statistical significance of survival curves using the log-rank test. We calculated the cytolytic activity (Cyt) as the geometric mean of *PRF1* and *GZMA* as previously described.^37^ We performed CIBERSORTx^38^ analysis on the RSEM gene expression of breast cancer patients to estimate the relative fraction of 22 immune cell types using 1000 permutations. We calculated the ssGSEA scores via the *GSVA* (v1.30.0) package^39^ using gene signatures collected from a previously described signature.^40^ We calculated in R the Spearman’s rank correlation coefficient between the median expression of 13 exosome signature genes and a single gene of interest.

## Results

### Establishing a single-cell method for quantifying exosome secretion

We sought to establish a method based on nanowell arrays for identifying the secretion of exosomes at the single-cell level. We have previously demonstrated that functionalized beads can serve as biosensors to enable the efficient capture of analytes from single cells within nanowell arrays.^28^ Accordingly, we wanted to investigate whether beads can serve to capture exosomes secreted by single cells within nanowell arrays.

The expression of transmembrane proteins, CD63 and CD81 on the surface of exosomes, has been widely used for isolation and detection of exosomes.^41^ We sought to compare the use of either a single marker (CD63) or two markers (CD63 and CD81) for the capture of exosomes. Accordingly, we isolated exosomes from GSC20 cancer cell line, using a standard ultracentrifugation procedure. Nanoparticle tracking analyses (NTA) confirmed that the exosomes had a median diameter of (132 ± 6) nm (Figure S1A). Quantitative analyses of the capture of purified exosomes onto either anti-CD63 or anti-CD81 antibody-coated beads demonstrated a specific increase in fluorescence when detected using a fluorescent anti-CD63 antibody (Ab). The beads coated with the anti-CD81 Ab showed lower background fluorescence in comparison to the anti-CD63-coated beads in the absence of exosomes (Figure S1B), which resulted in an increased area under the curve (AUC) (Figure S1C). Moving forward, we thus implemented the use of antibodies targeting CD81 (for capture) and CD63 (for detection).

To determine whether the immunoassay can capture exosomes secreted directly from cells, we modified the widely utilized transwell assay to harvest exosomes directly from cells.^42-44^ We chose to work with a pair of syngeneic, isogenic mouse breast cancer cell lines, with differing metastatic potential, 4T1 and 67NR. We incubated the non-metastatic 67NR mouse breast cells in the upper chamber with anti-CD81-coated beads in the lower chamber of a transwell assay for 48 hours (Figure S1D). The exosomes isolated using this procedure displayed the expected morphology and size as observed by Transmission Electron Microscopy (TEM) (Figure S1E). Collectively, these results suggest that the bead-based immunosandwich utilizing anti-CD81 and anti-CD63 Abs can be used to capture exosomes from cells and could be used for single-cell assays (Figure S1F).

To analyze the secretion of exosomes from single cells, we utilized a custom nanowell array containing 9216 wells, and co-incubated beads and breast cancer cells (Figure 1A and Figure S2). At two-hour intervals, we added the fluorescently tagged anti-CD63 Ab and imaged the entire nanowell array. As expected, individual cells demonstrated heterogeneity in exosome secretion (Figure 1B). We compared the frequency of single-cell secreting exosomes between 67NR, and the isogeneic, metastatic breast cancer cell line, 4T1. To quantify heterogeneity in secretory behavior and to estimate the relative rate of secretion between individual cells and across the cell lines, we restricted analyses to nanowells containing a single cell and a single bead. Within these nanowells, we used a combination of image segmentation, thresholding, and normalized fluorescent intensities to identify if individual cells were classified as secretors or non-secretors (detailed description in Figure S3). At each of the time points tested—two, four, and six hours— there was no difference in the frequency of single cells secreting exosomes, comparing 4T1 and 67NR (Figure 1C). Within all cells that secreted exosomes, we also compared the number of exosomes secreted per cell across 4T1 and 67NR single cells. Somewhat surprisingly, the non-metastatic cell line 67NR single cells secreted more exosomes per cell at each of the time points profiled (Figure 1D). Tracking the kinetics of exosome secretion in individual 4T1 and 67NR cells during a six-hour period revealed three major classifications for the cells: (1) a major subpopulation of cells which showed continuous secretion, (2) a subpopulation of cells that showed burst secretion at two hours, then subsequently stopped secreting, and (3) cells with burst secretion starting at four hours (Figure 1E). Taken together, these results established that while the overall frequencies of cells secreting exosomes are not necessarily different between metastatic and non-metastatic cell lines, individual cells showed differences in secretory behavior. These results also indicate that non-metastatic 67NR cells can secrete more exosomes per cell in comparison with metastatic 4T1 cells, a characteristic that cannot be observed by routine ultracentrifugation procedures.

**Figure 1.**
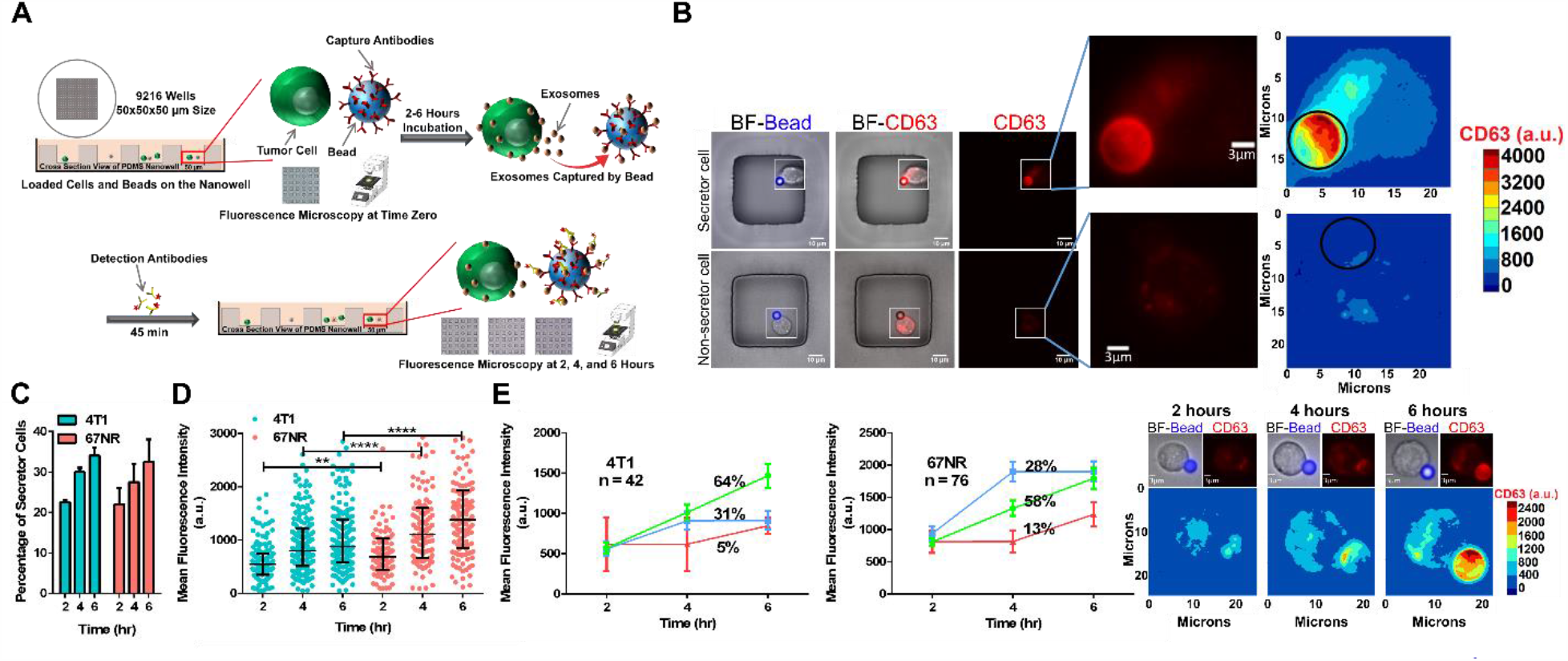
High-throughput single-cell assay for monitoring exosome secretion. **A**. The overall workflow of the single-cell assay. Cells and anti-CD81 conjugated beads are loaded on the nanowell and incubated for 2-6 hours. The entire array is incubated with fluorescently-labeled antibody against CD63 and imaged using microscopy. **B**. Representative images of individual nanowells containing 67NR cells with different exosome secretion capacity. The insets show single cells and the contour map of CD63 (exosome) intensity. **C**. Comparison of the frequency of exosome secreting cells between 67NR (non-metastatic) and 4T1 (metastatic) breast cancer cells. **D**. The rate of secretion of exosomes by 67NR cells is higher than that of 4T1 cells at two, four, and six hours. Each dot represents a single cell with the median and quantiles of CD63 (exosome) intensity shown across all cells. T-tests were used for comparison. A representative example from three independent repeats is shown. **E**. The kinetics of exosome secretion from single cells. The three subpopulations corresponding to (1) continuous secretion (green), (2) burst secretion at two hours (blue), and (3) burst secretion at four hours (red) are shown as trend lines (mean ± SEM). Representative images and contour maps of a single cell showing a continuous increase of CD63 intensity on the surface of the bead. Significance levels are shown as ** *p* < 0.001 and **** *p* < 0.00001.

### Single-cell RNA-sequencing illustrates that 67NR cells are enriched in exosome secretion pathways compared to 4T1 cells

To gain further mechanistic insights into the pathways that can support the increased exosome secretion capacity of 67NR cells in comparison to 4T1 cells, we performed single-cell RNA-sequencing (scRNA-seq). After data processing (see Methods), the final scRNA-seq dataset used for analyses had an average of 3,386 unique genes per cell and 35,604 transcripts (Figure S4A). Dimensionality reduction using t-Distributed Stochastic Neighbor Embedding (t-SNE) showed a clear separation between the cells comprising each cell line (Figure 2A). Hierarchical clustering indicates that a set of 1,647 differentially expressed genes (≥ 2-fold change) distinguishes the two cell types (Figure S4B). ScRNA-seq confirmed that a number of markers associated with epithelial-mesenchymal transition (EMT) including vimentin (*Vim*), fibronectin (*Fn1*), and Axl Receptor Tyrosine Kinase (*Axl*) were increased in 67NR cells (Figure S4C). In contrast, a number of matrix metalloproteinases associated with invasion, including *Mmp9* and *Mmp14*, were increased in 4T1 cells (Figure S4D).

**Figure 2.**
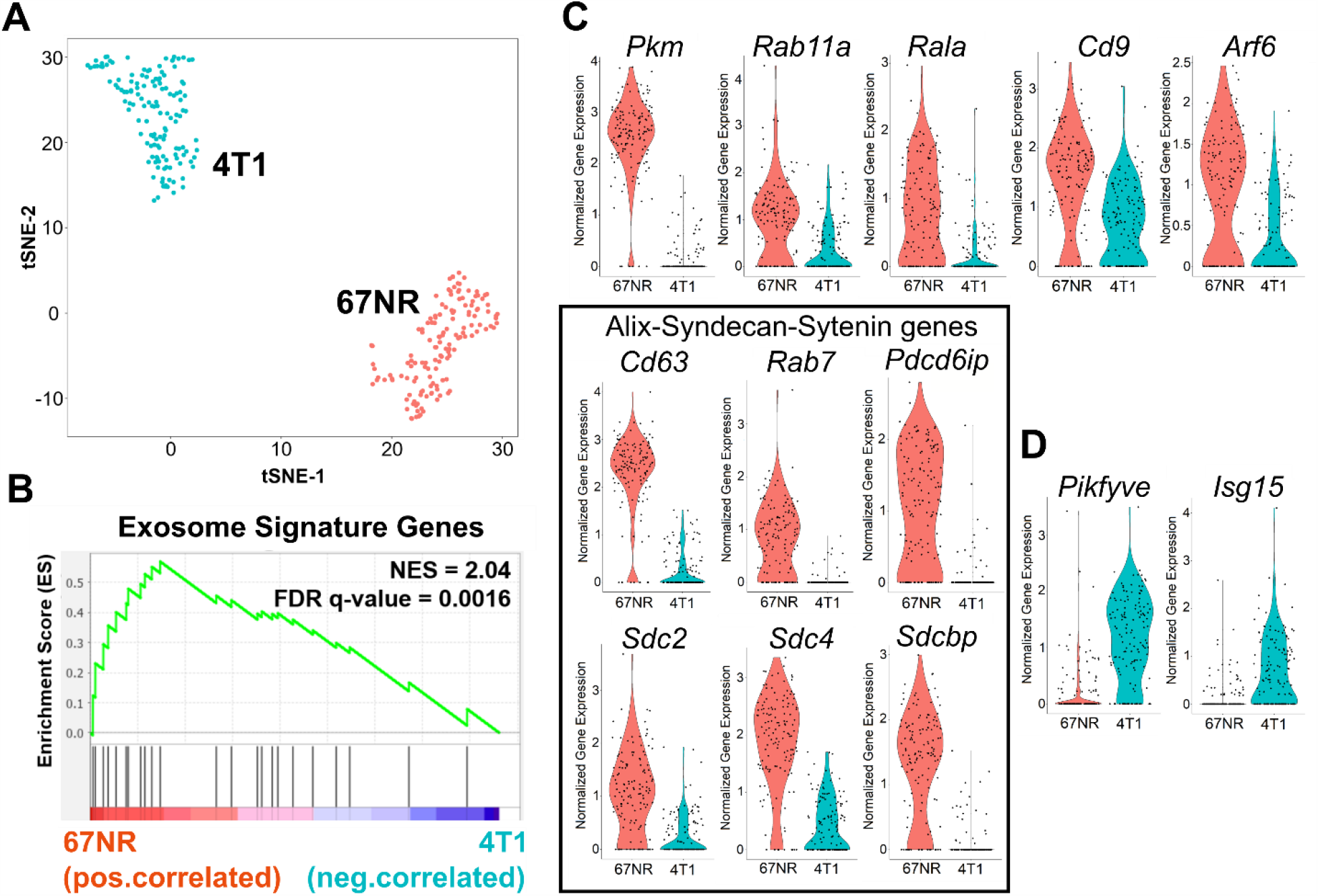
Comparison of 4T1 and 67NR cells by scRNA-seq. **A**. t-SNE plot of 67NR (non-metastatic) and 4T1 (metastatic) breast cancer cells clusters analyzed by scRNA-seq. **B**. Gene Set Enrichment Analysis (GSEA) analysis of the core exosome gene signature studies within 67NR cells compared to 4T1 cells. The core gene signature was based on a previously described set of genes known to be involved in exosome secretion (Table S1).^45^ **C**. Violin plots of the exosome-secretion genes that are different between 4T1 and 67NR. The black box highlights genes associated with ALIX-Syndecan-Sytenin pathway. **D**. Inhibitors of exosome secretion, *Pikfye* and *Isg15*, are enriched in 4T1 cells in comparison with 67NR cells.

To test the correlation between the functional single-cell exosome secretion assay and the transcriptional signatures, we established a core gene signature using a previously described set of genes known to be involved in exosome secretion (Table S1).^45^ Gene set enrichment analysis (GSEA) comparing 4T1 and 67NR confirmed that 67NR cells were positively correlated with exosome secretion signatures (Figure 2B, C). The core set of genes in the GSEA that showed high discrimination between 4T1 and 67NR cells mapped to the known ALIX-Syndecan-Sytenin pathway.^46^ The pathway genes consisting of tetraspanins (*Cd63*), *Rab7*, apoptosis-linked gene 2-interacting protein X (*Pdcd6ip*), syndecans (*Sdc2, Sdc4*), and syntenin (*Sdcbp*) were enriched in 67NR cells compared to 4T1 cells (Figure 2C). By contrast, two proteins that are known exosome secretion inhibitors, *Pikfyve* and *Isg15*, were significantly expressed in 4T1 cells but not in 67NR cells (Figure 2D).^47, 48^ We performed an independent verification of these results using reanalyzing population-level RNA-seq data on these same cell lines (GSE104765).^35^ These data also confirmed the higher expression of *Cd63, Rab7, Sdc2, Sdc3*, and *Sdcbp* in 67NR cells in comparison to 4T1 cells (Figure S4E). Collectively, these results from transcriptional profiling further advanced our findings that non-metastatic breast cancer cells can secrete more exosomes than metastatic breast cancer cells, and suggest that the ALIX-Syndecan-Sytenin pathway supports this function.

### Exosome secretion is an inheritable property during short-term culture of cancer cells

Our combined functional and transcriptional data illustrated that 67NR cells are proficient in exosome secretion. We next wanted to investigate the impact of exosome secretion on the functional properties of the 67NR tumor cells. We established a simple bioanalytical process to image cells secreting exosomes using nanowell arrays and microscopy, perform automated segmentation and identification of secretor and non-secretor cells, and use an automated micromanipulator to retrieve single cells to establish clonal cell lines (Figure 3A, Video S1). Since we want to ensure that long-term culture did not alter the properties of the cells, we grew the cells to no more than 24 population doublings. For the majority of the cells picked (20 out of 27), we were able to establish clonal cell lines, classified as secretor (cell population labeled as S, secretor, if the cell of origin was a secretor) and non-secretor (cell population labeled as NS, non-secretor, if the cell of origin was a non-secretor) [Figure 3B].

**Figure 3.**
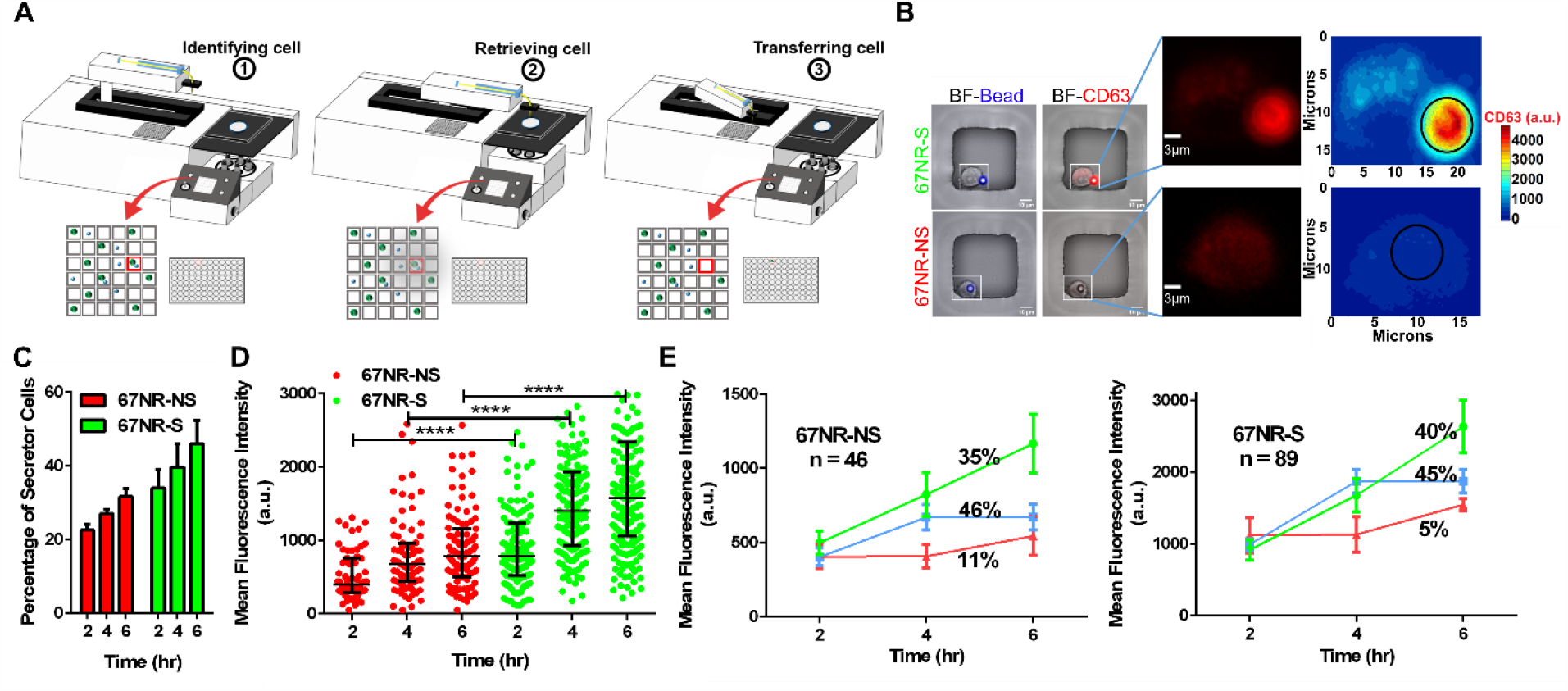
Establishment and validation of 67NR secretor (S) and non-secretor (NS) cell lines. **A**. Schematic of the overall workflow for the imaging and retrieval exosome secreting single cells with the aid of an automated micromanipulator. **B**. Representative images of 67NR secretor and non-secretor single cells before retrieval. The insets show single cells and the contour map of CD63 (exosome) intensity. **C**. Comparison of the frequency of cells secreting exosomes within expanded populations of 67NR-NS and 67NR-S cells. **D**. The rate of secretion of exosomes by cells within the 67NR-S population is higher than the rate of secretion by cells within the 67NR-NS population at two, four, and six hours. Each dot represents a single cell with the median and quantiles of CD63 (exosome) intensity shown in all cells. T-tests were used for comparison. A single representative experiment from four independent repeats is shown. **E**. The kinetics of exosome secretion from individual cells that comprise the 67NR-NS and 67NR-S populations. The three subpopulations corresponding to (1) continuous secretion (green), (2) burst secretion at two hours (blue), and (3) burst secretion at four hours (red) are shown as trend lines (mean ± SEM). Significance levels are shown as **** *p* < 0.00001.

We tested the ability of single cells derived from these expanded populations to secrete exosomes using the single-cell assay. Consistently, across all six cell lines tested (three secretor lines and three non-secretor lines), the frequency of single cells secreting exosomes was higher among the 67NR-S cell lines in comparison to the 67NR-NS cell lines (Figure 3C). Within all cells that secreted exosomes, comparisons of the number of exosomes secreted per single cell as a function of time (two, four, and six hours) confirmed that the 67NR-S cell lines were composed of individual cells with high rates of exosome secretion (Figure 3D). The numbers of exosomes secreted by single cells from the 67NR-NS cell lines and the 67NR-S cell lines were, respectively, lower and higher than the parental unsorted 67NR cell line (Figure S5). Kinetic analyses of the dynamics of exosome secretion in these cell lines revealed two dominant subpopulations: (a) continuous secretors and (b) cells with burst secretion that stopped secretion after 4 hours (Figure 3E). Taken together, these results establish that the secretion of exosomes is inheritable during cell division, and this allowed us to investigate the functional consequences of these exosome secreting cell lines.

### Secretion of exosomes prevents the tumor formation in non-metastatic cell lines

Since the expanded cell populations preserved the exosome secretion property of the cell of origin, we investigated *in vitro* functions of the 67NR-S and 67NR-NS cell lines. Phase-contrast microscopy revealed differences in the morphology with 67NR-S cells being more elongated than 67NR-NS cells (Figure 4A). Migration is a key characteristic of cancer cells essential for metastasis. To test the migratory behavior of the 67NR-S and 67NR-NS cell lines, we performed a scratch wound assay.^49^ 67NR-S cells were significantly more migratory than 67NR-NS cells (Figure 4B). To test the tumorigenicity potential of these cell lines, we used a soft agar formation assay. 67NR-S cells formed 2-fold more colonies than the 67NR-NS cells in soft agar suspension cultures (Figure 4C). These *in vitro* data illustrate that the 67NR-S cells were more migratory and had enhanced tumorigenicity potential compared to 67NR-NS cells.

**Figure 4.**
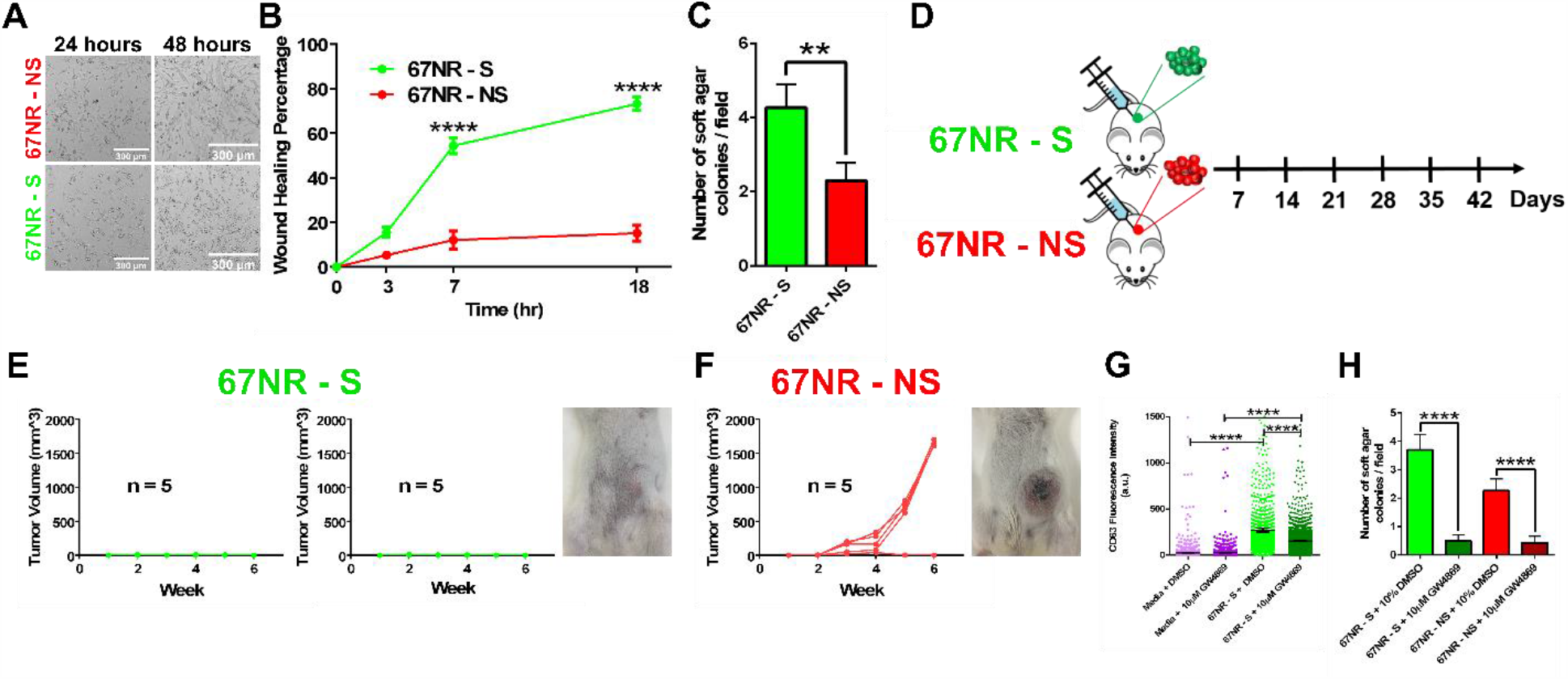
Assessing the functionality of 67NR-NS and 67NR-S cells. **A**. The morphology of 67NR-NS and 67NR-S cell populations recorded using phase-contrast microscopy. **B**. Wound healing assays illustrating the migration of 67NR-S clonal cell populations in comparison with 67NR-NS clonal cell populations (mean ± SEM). A two-way ANOVA test was used (n= 6 for each cell line). **C**. 67NR-S cell populations have a higher capacity to form colonies in comparison with 67NR-NS cell populations (mean ± SEM). The Mann Whitney t-test was used for comparison. **D**. The design of mice experiments for comparing the efficacy of tumor formation by 67NR-S and 67NR-NS cell lines. **E**. Tumor growth monitoring of BALB/c mice injected with 67NR-S clones (two clonal cell populations, five mice each). A representative image of a single mouse is shown. **F**. Tumor growth monitoring of BALB/c mice injected with 67NR-NS clone (single clonal cell population, five mice). A representative image of a single mouse is shown. **G**. Inhibition of exosome secretion within 67NR-S clonal cell populations by 10 µM GW4869 compared to DMSO control using a 48-hour transwell assay. Media containing either 10 µM GW4869 or DMSO were used as a negative control. Each dot represents CD63 (exosomes) intensity on a single bead (mean ± SEM). The Mann Whitney t-test was used for comparison. **H**. Inhibition of colony formation in 67NR-S and 67NR-NS clonal cell populations upon treatment with 10 µM GW4869 (mean ± SEM). The Mann Whitney t-test was used for comparison. Significance levels are shown as ** *p* < 0.01, and **** *p* < 0.0001.

We have utilized syngeneic models to be able to understand the impact of exosomes on both intrinsic growth potentials of the tumor and the impact of the host immune system. Parental 67NR cells are non-metastatic cells with a heterogeneous population and form primary tumors upon injection into mice. To determine the *in vivo* relevance of exosome secretion, we injected two 67NR-S and one 67NR-NS cell lines into the mammary fat pad of BALB/c mice and monitored the tumor growth for six weeks (Figure 4D). None of the mice that received the 67NR-S cells developed tumors (Figure 4E). By comparison, however, 80% of the mice that received 67NR-NS cells formed large tumors by week six (Figure 4F). Taken together, these results illustrate that despite having high tumorigenicity and migratory potential *in vitro*, the 67NR-S cells are rejected *in vivo*.

To directly link exosome secretion to the rejection of tumors *in vivo*, we investigated the use of GW4896, a chemical inhibitor of exosome biogenesis. Treatment of 67NR-S cells with GW4896 significantly inhibited exosome secretion when profiled using the transwell exosome capture assay (Figure 4G). Unfortunately, however, the treatment of 67NR-S cells with GW4896 almost completely abolished colony formation in a soft agar assay (Figure 4H), precluding its use *in vivo*. Collectively, studies with these non-metastatic breast cancer cells demonstrated that despite enhanced tumor-forming potential *in vitro*, exosome secreting cell lines are rejected *in vivo* presumably due to the host immune system.

### Secretion of exosomes improves the survival in non-metastatic breast cancer patients

Based on the mice data, we sought to directly understand the impact of exosome secretion and the link to the immune system within human patients with breast cancer. We analyzed the correlation between gene expression and survival of non-metastatic breast cancer patients available within The Cancer Genomic Atlas (TCGA). Since our single-cell method utilizes CD63 and CD81 to detect the exosomes, we first compared the survival of patients with higher and lower expression of these markers. Since there was no difference in survival of patients stratified by CD63 or CD81, we conclude that these single markers are necessary but not sufficient to identify a complex property like exosome secretion (Figure 5A, Figure S6A).

**Figure 5.**
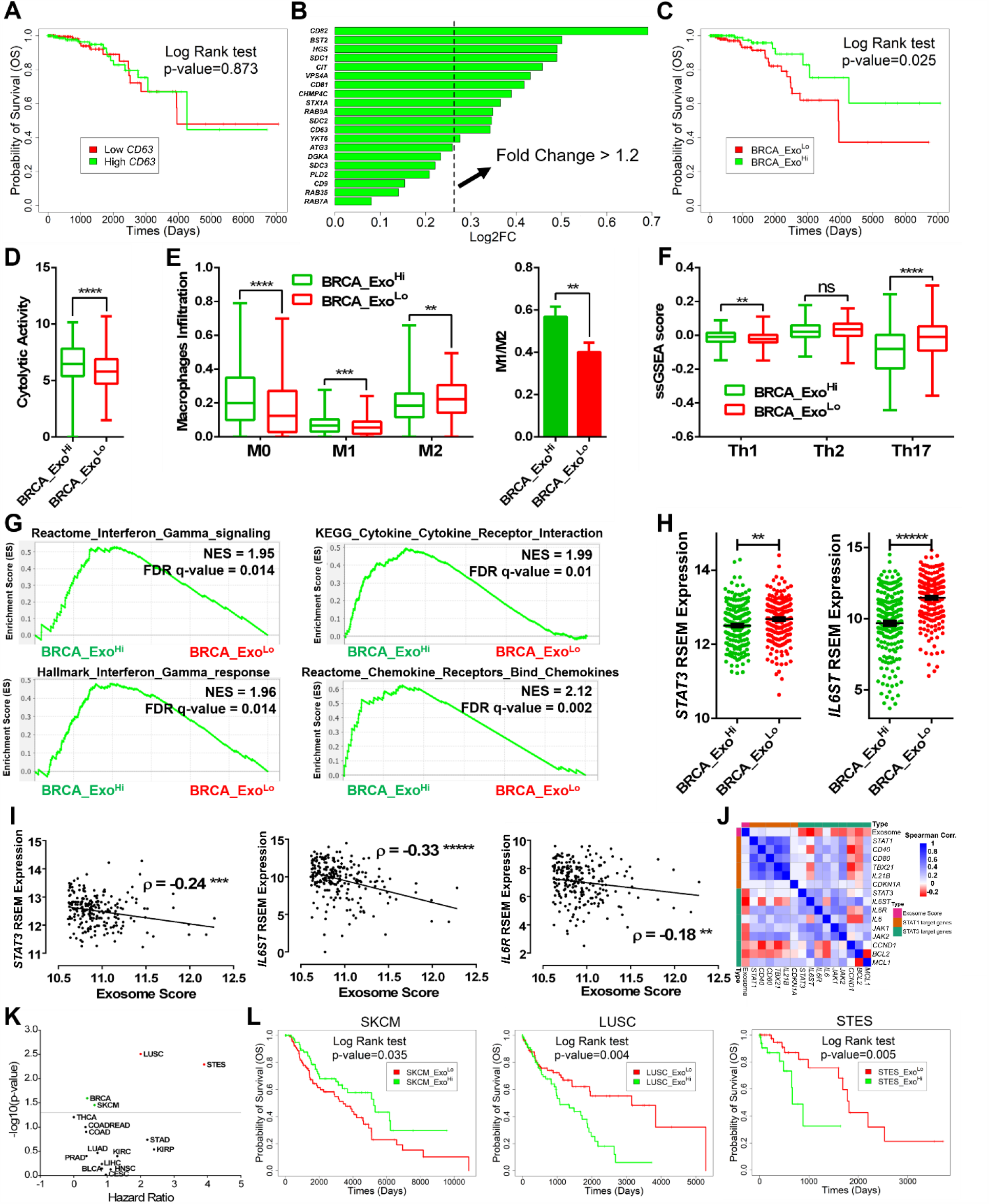
Increased exosome secretion is correlated with better survival in non-metastatic breast cancer patients. **A**. The overall survival of non-metastatic breast cancer patients (N0 and M0 in TNM staging system) divided by median *CD63* expression. **B**. Enrichment of the genes associated with exosome secretion in the BRCA_Exo^Hi^ patients. The 13 genes with fold change > 1.2 were selected as the exosome gene signature. **C**. Differences in the survival of non-metastatic breast cancer patients stratified by the median expression of 13 exosome gene signature. **D**. Cytolytic activity score of non-metastatic breast tumors comparing the BRCA_Exo^Hi^ and BRCA_Exo^Lo^ patients. The median and quantiles of the scores are shown. **E**. Macrophage infiltration scores for the BRCA_Exo^Hi^ and BRCA_Exo^Lo^ tumors determined by CIBERSORTx. The median and quantiles of the infiltration fraction are shown. The ratio of M1/M2 macrophages within these tumors is also shown (mean ± SEM). **F**. The immune score of T helper cells for BRCA_Exo^Hi^ and BRCA_Exo^Lo^ tumors identified using single sample Gene Set Enrichment Analysis (ssGSEA). The median and quantiles of the infiltration percentage are shown. **G**. GSEA of interferon-gamma, cytokines/chemokines receptor interaction pathways comparing BRCA_Exo^Hi^ and BRCA_Exo^Lo^ tumors. **H**. Normalized expression of *STAT3* and *IL6ST* in BRCA_Exo^Hi^ and BRCA_Exo^Lo^ tumors (mean ± SEM). **I**. The anti-correlation of *STAT3, IL6R*, and *IL6ST* with exosome score within non-metastatic breast cancer patients (Spearman correlation). **J**. Spearman correlation between genes of the *STAT1* and *STAT3* pathways and the genes associated with exosome secretion within non-metastatic breast cancer patients. **K**. Volcano plot of overall survival of pan-cancers divided by the median expression of exosome gene signatures. **L**. Overall survival of non-metastatic SKCM, LUSC and STES patients divided by median expression of exosome gene signatures. For E, F, G, and I, two-tailed t-test was used. Significance levels are shown as * *p* < 0.05, ** *p* < 0.01, *** *p* < 0.001, **** *p* < 0.0001, ***** *p* < 0.00001.

To identify signatures of exosome secretion, we applied unsupervised hierarchal clustering (no gene selection) to stratify non-metastatic breast cancer patients into two groups with 182 and 268 patients each (Figure S6B). A set of 13 genes related to exosome secretion were identified as being differentially expressed between these two groups (Figure 5B). We therefore utilized the median expression of this 13-gene cluster to stratify patient tumors as exosome high (BRCA_Exo^Hi^) and low (BRCA_Exo^Lo^). Consistent with our scRNA-seq data on 4T1 and 67NR cells, the expression of genes in ALIX-Syndecan-Sytenin pathway was elevated in BRCA_Exo^Hi^patients in comparison to the BRCA_Exo^Lo^ patients (Figure S6C). The overall survival was significantly higher for BRCA_Exo^Hi^ patients in comparison to the BRCA_Exo^Lo^ patients (median survival not reached vs. 10.8 years, HR: 0.4, 95% CI: 0.18-0.92), consistent with our findings in mice that non-metastatic cells secreting exosomes do not form tumors (Figure 5C).

To identify if immune cell infiltration is associated with improved overall survival observed in patients with higher expression of exosomes, we used the previously published cytolytic score (based on the expression of *GZMA* and *PRF1*) as an *in silico* metric of immune cell cytolytic activity.^37^ The cytolytic activity was significantly elevated in the BRCA_Exo^Hi^ cohort compared to the BRCA_Exo^Lo^ cohort (Figure 5D). To identify the immune cell type that was responsible for this signature, we used the normalized gene expression data to quantify the relative frequencies of the 22 different immune cell types using the CIBERSORTx algorithm. CD8 T cells were not significantly different between the two clusters (Figure S6D). The difference in cytolytic activity was reflected with significant differences in macrophage subsets: a higher frequency of M0 and pro-inflammatory M1 macrophages, and a decreased frequency of anti-inflammatory M2 macrophages were observed in the tumors of BRCA_Exo^Hi^ patients compared to the BRCA_Exo^Lo^ patients (Figure 5E). Similar to the macrophages, the frequency of intratumoral memory CD4 T cells was also significantly different between BRCA_Exo^Hi^ and BRCA_Exo^Lo^ patients. We utilized signatures of helper T cells within the previously described Immunome signature set,^40^ to identify that Th1 cells were significantly increased, and Th17 cells were significantly decreased in the BRCA_Exo^Hi^ patients compared to the BRCA_Exo^Lo^ patients (Figure 5F). Collectively, these results showed that tumors in BRCA_Exo^Hi^ patients harbor M0/M1 macrophages and Th1 cells.

We utilized GSEA to identify soluble mediators of the immune cell polarization within the tumor microenvironment of these patients. Not surprisingly, several pathways associated with chemokine/cytokine receptor interactions were enriched in BRCA_Exo^Hi^ tumors (Figure 5G). Consistent with the high frequency of Th1 cells, interferon-gamma (IFN-γ) signaling was significantly elevated within the exosome high tumors (Figure S6E). It is well known that the priming of macrophages in the presence of IFN-γ leads to the differentiation of pro-inflammatory M1 macrophages and downregulation of the IL6 signaling pathway. Although the expression of the IL6 receptor (*IL6R*) was not different, the expression of IL6 signal transducer (*IL6ST*) and the downstream signal transducer and activator of transcription 4 (*STAT3*) were significantly decreased in the BRCA_Exo^Hi^ tumors compared to the BRCA_Exo^Lo^ tumors (Figure 5H). We also utilized the median expression score of the 13 exosome signature genes to confirm a significant inverse correlation between the exosome signature and *IL6R, IL6ST*, and *STAT3* within this entire cohort of patients (Figures 5I and 5J). Taken together, the secretion of exosomes likely influences the infiltration of Th1 cells and a skewed ratio of M1/M2 macrophages through the crosstalk between IFN-γ and IL6/STAT3 pathways.

We investigated the utility of the exosome secretion signature and its association with patient survival across pan-cancer datasets within the TCGA. Similar to breast cancer, exosome secretion signatures were associated with improved overall survival in melanoma (SKCM) patients (14.3 vs. 9.4 years, HR: 0.62, 95% CI: 0.39-0.97) [Figure 5K]. By contrast, in both lung squamous cell carcinoma (LUSC) and stomach and esophageal carcinoma (STES), exosome secretion signature was associated with worse overall survival for patients (Figure 5L).

## Discussion

Exosomes and EVs derived from tumors serve as long-distance messengers and hence play important roles in the metastatic cascade.^50, 51^ Exosomes derived from metastatic cells have been shown to participate in a broad range of functions: remodel the extracellular matrix and the transformation of fibroblasts, promote angiogenesis, prepare the pre-metastatic niche, and alter the nature of the tumor microenvironment.^51-53^ By contrast, the exosomes released by non-metastatic tumors has been relatively understudied. In both metastatic and non-metastatic tumors, studies that have aimed to investigate the role of exosomes have utilized exosomes purified from large numbers of cancer cells isolated from cell culture. This approach masks the heterogeneity in the secretion of the different cells that comprise the population. Comprehensive characterization of the heterogeneity of exosome secretion between the single cells within the same population has been restricted to only a few reports, and even in these studies, the ability to isolate and propagate cells with differences in exosome secretion capabilities is lacking.^24, 54^ We developed and validated a platform based on nanowell arrays for directly profiling exosome secretion from single cells and used these to establish cells derived from a clinically relevant mouse breast cancer model that have significant differences in the rate of exosome secretion. Our studies show that, surprisingly, the non-metastatic cell line, 67NR secretes more exosome per cell than its isogenic, metastatic counterpart, 4T1. Although prior studies from each of these cell lines have demonstrated that exosomes derived from 4T1 can facilitate metastasis, these studies utilized supraphysiological concentrations of purified exosomes.^55^ Our results are consistent with studies in melanoma that showed that purified exosomes from poorly metastatic melanoma cells could inhibit metastasis.^56^ ScRNA-seq suggested that the ALIX-Syndecan-Sytenin pathway known to be important for exosome secretion was enriched in 67NR cells compared to 4T1 cells.^46^ *In vitro* functional studies based on 67NR-S (high secretor) and 67NR-NS (low secretor) cells illustrated that the 67NR-S cells are more migratory and have enhanced tumorigenic potential; however, they are deficient at tumor formation *in vivo*.

To explore the relevance of our results in human breast cancers, we analyzed the signatures of exosomes with non-metastatic breast cancer patients within the TCGA. Our results demonstrate that patients with signatures of high exosome secretion (including the ALIX-Syndecan-Sytenin pathway) have improved survival compared to patients with signatures of low exosome secretion. To quantify if immune cells can help explain this difference in survival between the two cohorts of patients, we quantified the cellular composition in terms of the 22 subtypes of immune cells using the CIBERSORTx algorithm.^38^ The cytolytic score, an *in silico* metric of inflammation, was significantly increased in tumors with high exosome secretion compared to tumors with low exosome secretion.^37^ Surprisingly, the cytolytic score was not reflected by the high abundance of CD8 T cells but correlated with Th1 cells (secretion of IFN-γ) and M1 macrophages (suppression of IL6ST/STAT3 pathway). These results are consistent with studies using purified exosomes that revealed that breast cancer-derived exosomes alter macrophage polarization via IL6ST/STAT3 signaling.^57^

Our platform has direct utility in single-cell studies of profiling the link between exosomes and function. In the current report, we have defined exosome secretion based only on abundances of CD63/CD81. This definition only marks a subset of all exosomes, but the assay can be easily modified to include additional markers including CD9 and EpCAM.^58^ Second, the ability to isolate cells based on differences in exosome secretion can be utilized to perform scRNA-seq on the retrieved cells directly. This method will have great utility to map the molecular players in the exosome secretion cascade directly. Furthermore, based on the differentially expressed transcripts, it will be possible to infer the proteins that are likely enriched in the exosomes secreted by these single cells. Third, the establishment of cell lines with differences in exosome secretion among metastatic cells will help map the functional impact of exosome secretion and their role in the biology of metastasis. We anticipate that our method can be broadly utilized to map the functional consequences of exosome secretion at the single-cell level.

## Supporting information

Single cell retrieval

## Acknowledgments

We would like to acknowledge the MDACC Flow Cytometry and Cellular Imaging Core facility for the FACS sorting (NCI P30CA16672) and MDACC High Resolution Electron Microscopy Facility for TEM microscopy (CCSG grant NIH P30CA016672). We would like to thank Ali Rezvan for the fabrication of the nanowell arrays.

## Declaration of Interest Statement

UH has filed a provisional based on some of the technologies described in this manuscript.

## Author Contributions

N.V. and M.F. designed the study. X.A. fabricated PDMS nanowell arrays. M.F., R.J., M.M.P., and X.A. performed *in vivo* study. M.F., and J.T.A analyzed single-cell RNA-seq. M.F. performed TCGA analysis. K.G. performed exosome isolation. S.A.M., and N.V. supervised the study.

## Supplemental Materials

The supplemental materials are available free of charge. Additional figures and tables (PDF) Single cell retrieval (AVI)

## Funding

This work was supported in part by the NIH under grant U01AI148118; CPRIT under grant RP180466; MRA Established Investigator Award under grant 509800; NSF under grant 1705464; CDMRP under grant CA160591; and Owens foundation to N.V., Cancer Prevention Research Institute of Texas (CPRIT) Multi-Investigator Research Award (MIRA) under grant RP160710 to S.A.M.

## Notes

### Competing Interest Statement

NV is the founder and CSO of CellChorus. UH has filed patent applications based on some of the technologies described in this study.

