## Supplementary Data for "Functional single-cell profiling identifies that exosomes are associated with increased immune cell infiltration in non-metastatic breast cancer"

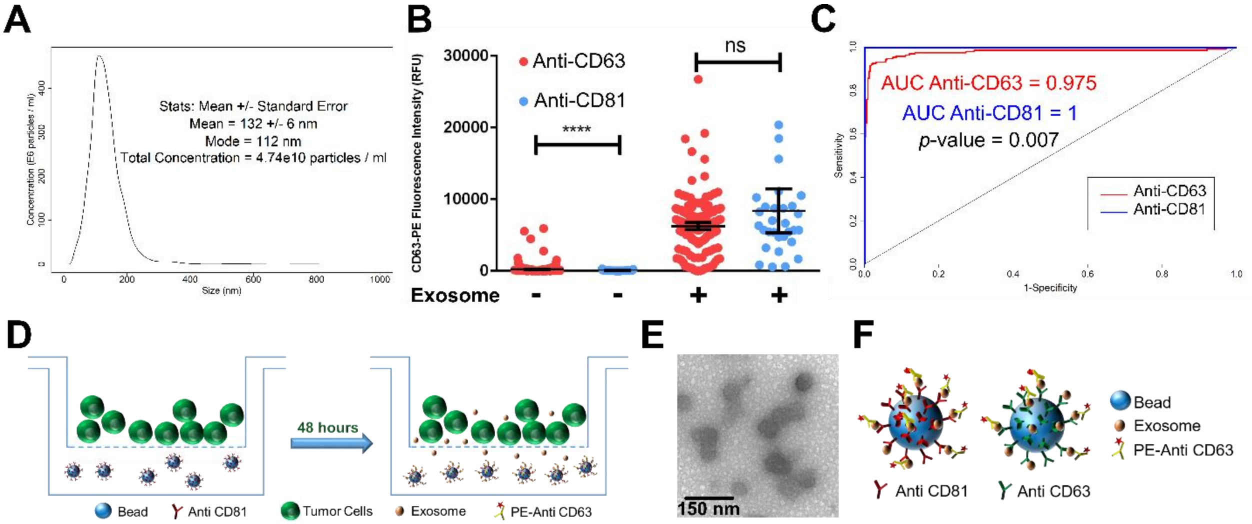


**Figure S1. Optimization of exosome capture using bead based immunoassay.**

A. Nanosight analysis of exosomes isolated from GSC20 cells.

B. CD63 (exosome) intensity of beads functionalized with either anti-CD63 or anti-CD81 for the capture of exosomes. PBS was used as negative control. Each dot represent a single bead. Two-tailed t-test was applied.

C. The use of the anti-CD81 as the capture antibody leads to accurate detection of exosomes. DeLong's test was applied for two ROC curves.

D. Overall workflow of transwell assay for capturing exosomes for TEM visualization.

E. TEM images of exosomes isolated using the transwell assay.

F. Overall representative schematic of immunoassay showing higher efficiency of anti CD81 for capturing exosomes. Significance levels are shown as **** $p < 0.0001$.

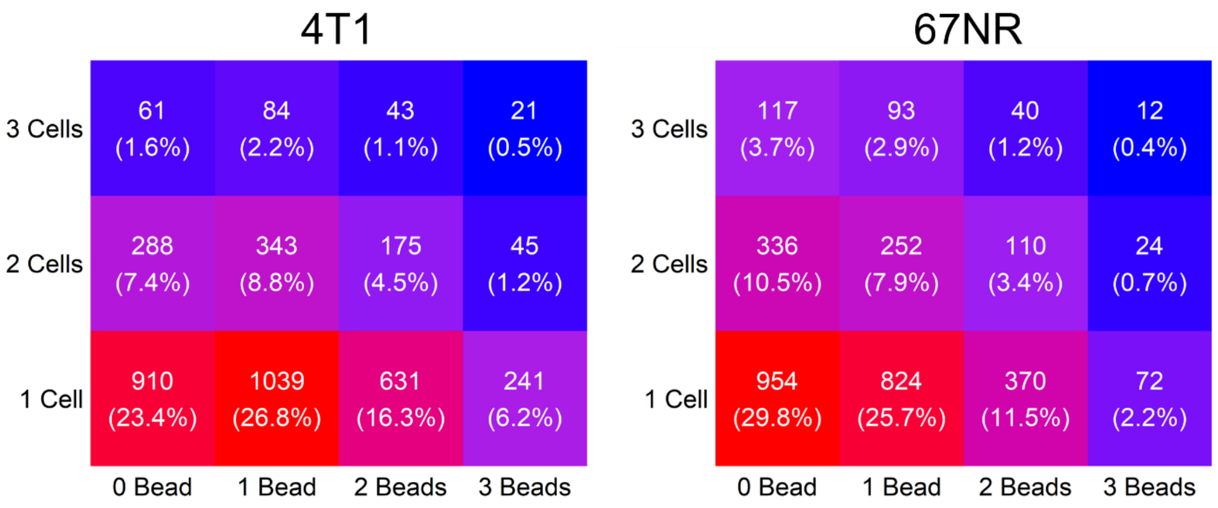


**Figure S2.** Distribution of bead:cell ratio within nanowell arrays. Two representative experiments are shown.

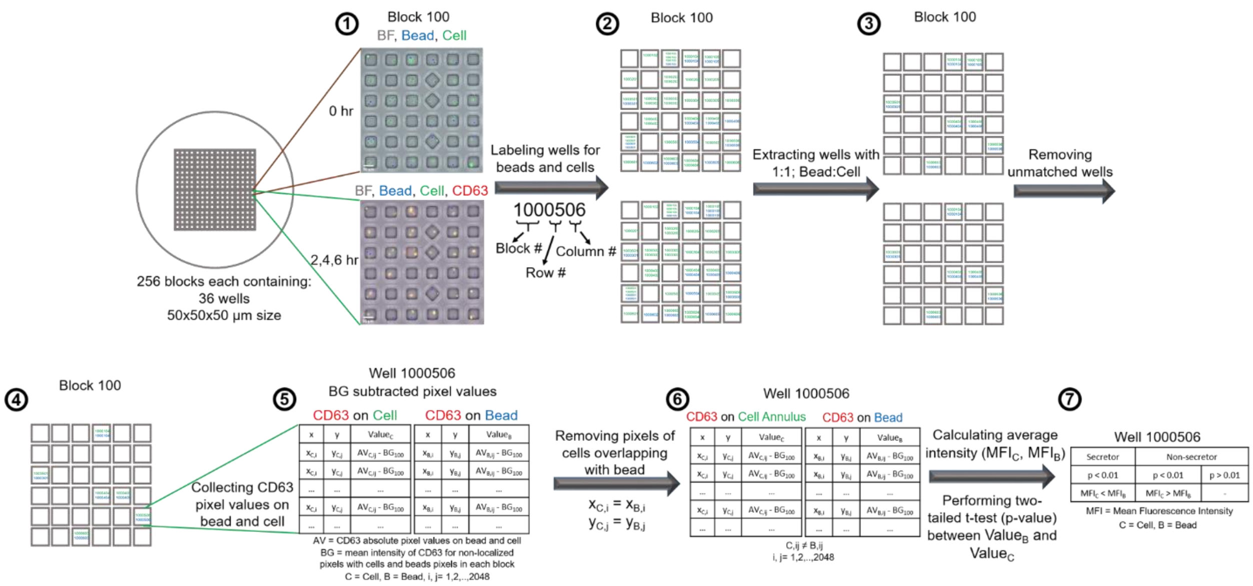


**Figure S3. Overall workflow of the automate image-analysis for identification of secretor and non-secretor cells.**

1. TIFF images of 256 field of views (blocks) containing 36 wells were extracted and merged for the initial time point (0 hour) and the detection time points (2, 4, 6 hours). Block 100 is shown as an example for workflow.

2. The numbers of beads and cells in each well were identified for each block at all the time points.

3. Wells with a single bead and a single cell were identified.

4. The wells which didn't maintained bead:cell = 1:1 ratio in entire experiment were excluded from the further analysis.

5. The CD63 pixel values on the surface of cell and bead were collected. These values were corrected using background subtractions. To calculate background intensity, average of CD63 pixel values not localized on the beads and cells in the entire block was calculated. Well 1000506 is shown as an example for workflow.

6. The overlap pixels between bead and cell were removed from pixel value sets of cell. This created an annulus shape for the pixels on the surface of cell.
7. Two-tailed t-test was applied on two set of pixel values, on bead and on the cell annulus, to identify the secretor cells in which the CD63 intensity for beads was significantly ($p < 0.01$) higher than cell annulus intensity.

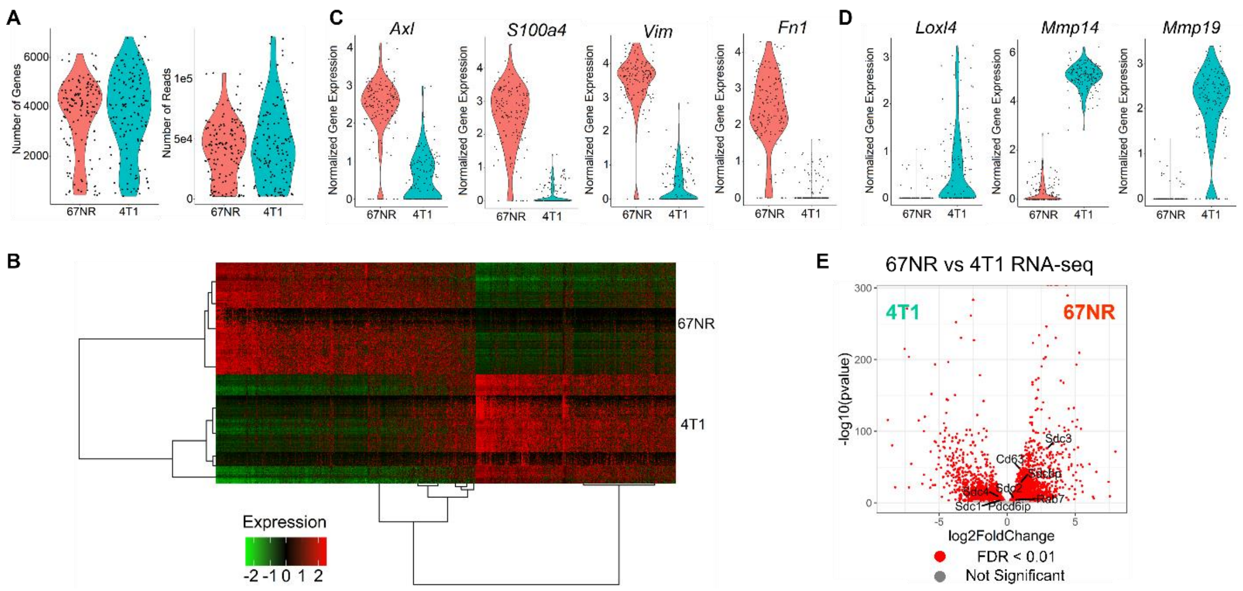


**Figure S4. Transcriptome comparison of 67NR (non-metastatic) and 4T1 (metastatic) cells by single-cell and bulk RNA-seq.**

A. Number of genes and reads detected in scRNA-seq for 67NR and 4T1 cells.

B. Heat map comparing the expression of the differentially expressed genes in 67NR and 4T1 cells.

C. Violin plot comparing the expression of mesenchymal cell transcripts.

D. Violin plot comparing the expression of epithelial cell transcripts.

E. Expression of genes associate with Alix-Syndecan-Sytenin pathway in 67NR cells in comparison with 4T1 cells based on the bulk RNA sequencing.

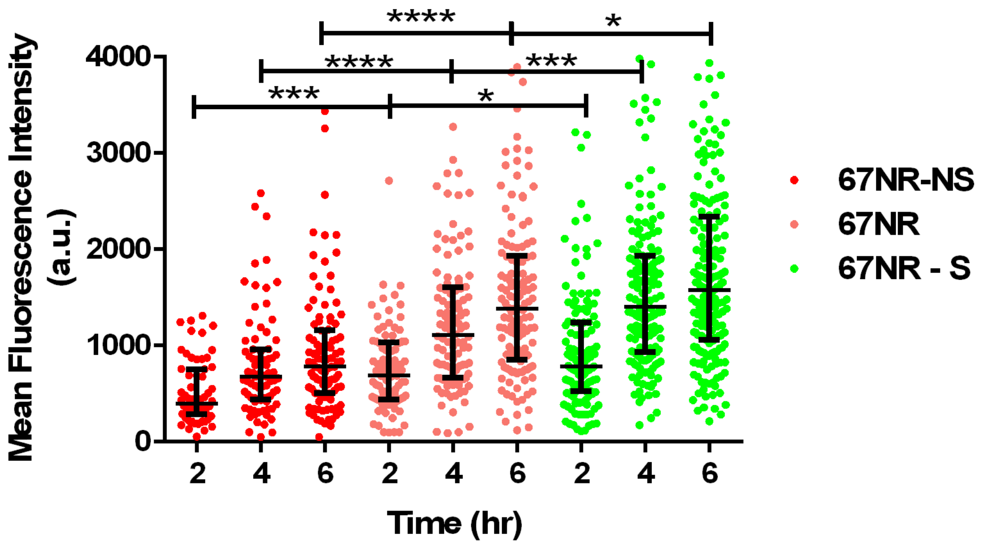


**Figure S5.** Higher secretion rate for 67NR-S cells and lower secretion rate for 67NR-NS cell in comparison to parental 67NR cells. Each dot represents a single cell with the median and quantiles of CD63 (exosome) intensity shown over all cells. T-tests were used for comparison. Significance levels are shown as * $p < 0.01$, ** $p < 0.001$, *** $p < 0.0001$, **** $p < 0.00001$, ***** $p < 0.000001$.

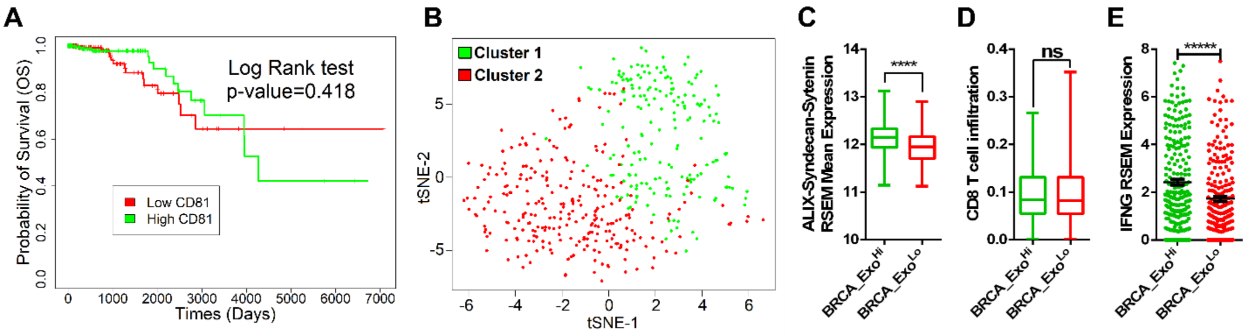


**Figure S6. Exosome secretion effects on TCGA patients.**

A. Overall survival of non-metastatic breast cancer patients (N0 and M0 in TNM staging system) divided by CD81 median expression.

B. T-SNE plot of the clusters identified by unsupervised hierarchal clustering of non-metastatic breast cancer patients.

C. The average expression of genes in ALIX-Syndecam-Syntenin pathway (*CD63*, *SDC1*, *SDC2*, *SDC3*, *SDC4*, *SDCBP*, *RAB7A*, *PDCD6IP*) within BRCA_Exo$^{Hi}$ and BRCA_Exo$^{Lo}$ patients. The median and quantiles are shown.

D. Infiltration of CD8 T cells within BRCA_Exo$^{Hi}$ and BRCA_Exo$^{Lo}$ patients. The median and quantiles of the infiltration fraction are shown.

E. High expression of *IFNG* in BRCA_Exo$^{Hi}$ patients in comparison to BRCA_Exo$^{Lo}$ patients (mean ± SEM). Two-tailed t-test was used. Significance levels are shown as **** $p < 0.0001$.

**Table S1.** Gene signature associated with exosome secretion.

| | Mouse symbol | Human Symbol | Ref. | | Mouse symbol | Human Symbol | Ref. |
|---|---|---|---|---|---|---|---|
| 1 | *Hgs* | *HGS* | [1-4] | 23 | *Pkm* | *PKM* | [5] |
| 2 | *Stam* | *STAM* | [1] | 24 | *Snap23* | *SNAP23* | [5] |
| 3 | *Tsg101* | *TSG101* | [1] | 25 | *Rala* | *RALA* | [6] |
| 4 | *Chmp4c* | *CHMP4C* | [1] | 26 | *Ralb* | *RALB* | [6] |
| 5 | *Pdcd6ip* | *PDCD6IP* | [1, 7] | 27 | *Rab2b* | *RAB2B* | [8] |
| 6 | *Vta1* | *VTA1* | [1] | 28 | *Rab5a* | *RAB5A* | [8] |
| 7 | *Vps4a* | *VPS4A* | [1] | 29 | *Rab9* | *RAB9A* | [8] |
| 8 | *Chmp4c* | *CHMP4C* | [1] | 30 | *Rab7* | *RAB7A* | [7, 9] |
| 9 | *Sdcbp* | *SDCBP* | [7] | 31 | *Rab11a* | *RAB11A* | [10, 11] |
| 10 | *Sdc1* | *SDC1* | [7] | 32 | *Rab27a* | *RAB27A* | [4, 8, 12-14] |
| 11 | *Sdc2* | *SDC2* | [7] | 33 | *Rab27b* | *RAB27B* | [8, 9] |
| 12 | *Sdc3* | *SDC3* | [7] | 34 | *Rab35* | *RAB35* | [15, 16] |
| 13 | *Sdc4* | *SDC4* | [7] | 35 | *Cit* | *CIT* | [17] |
| 14 | *Cd9* | *CD9* | [18] | 36 | *Cttn* | *CTTN* | [19] |
| 15 | *Cd82* | *CD82* | [18] | 37 | *Smpd3* | *SMPD3* | [20-22] |
| 16 | *Cd63* | *CD63* | [23] | 38 | *Dgka* | *DGKA* | [24] |
| 17 | *Lmp1* | *LMP1* | [25] | 39 | *Pld2* | *PLD2* | [26, 27] |
| 18 | *Tspan8* | *TSPAN8* | [28] | 40 | *Arf6* | *ARF6* | [27] |
| 19 | *Syt7* | *SYT7* | [4] | 41 | *Bst2* | *BST2* | [29, 30] |
| 20 | *Vamp7* | *VAMP7* | [31] | 42 | *Atg12* | *ATG12* | [32] |
| 21 | *Ykt6* | *YKT6* | [3, 33] | 43 | *Atg3* | *ATG3* | [32] |
| 22 | *Stx1a* | *STX1A* | [11] | | | | |
